# Modeling Macroscopic Currents of Ion Channels

**DOI:** 10.1101/2021.11.14.468518

**Authors:** Di Wu

## Abstract

Ion-channel functions are often studied by the current-voltage relation, which is commonly fitted by the Boltzmann equation, a powerful model widely used nowadays. However, the Boltzmann model is restricted to a two-state ion-permeation process. Here we present an improved model that comprises a flexible number of states and incorporates both the single-channel conductance and the open-channel probability. Employing the channel properties derived from the single-channel recording experiments, the proposed model is able to describe various current-voltage relations, especially the reversal ion-permeation curves showing the inward- and outward-rectifications. We demonstrate the applicability of the proposed model using the published patch-clamp data of BK and MthK potassium channels, and discuss the similarity of the two channels based on the model studies.

## Introduction

Under particular environmental stimuli, certain types of ion channels open and transmit specific ions across biological membranes that evoke subsequent actions of more types of ion channels cooperating at the proper time, a process known to control or influence many physiological and neural activities (1), including generating the action potentials, releasing hormones and neurotransmitters, regulating skeletal and cardiac muscle contraction, controlling cell volume, affecting cell proliferation, fertilization and cell death, etc. Ion channel functions are usually studied by the patchclamp recording methods (2, 3). In the voltage-clamp recordings, the current traces responding to various voltage pulses are recorded, and the current is conventionally calculated by I = *N*P_o_*γ*(*V* – *V*_rev_), where *N* is the number of channels in the patch, P_o_ is the channel open probability, *γ* is the unitary single-channel conductance, *V* is the membrane potential (controlled by the test voltage in the voltage-clamp experiments), and *V*_rev_ is the reversal potential of the permeant ions. The channel open probability P_o_ of a single channel, or *N*P_o_ of all channels, can vary with voltage *V*, which is usually fitted by the Boltzmann equation derived from the probability theory proposed by Hodgkin and Huxley (4). Various versions of Boltzmann equation exist, differing in only the expressions of the exponent parameters, but they all represent the same theory that the probability of the open state relative to the closed state is proportional to a Boltzmann factor. Here we use the expression: P_O_ = 1/(1 + exp(*q*(*V* – *V*_1/2_)/*k_B_T*)), where *V* represents the membrane potential, *V*_1/2_ is the half activation potential, *q* is the electric-charge change associated with the activation of the channel, *k*_B_ is Boltzmann’s constant, and *T* is the absolute temperature. Over decades of experience, Boltzmann equation proves to be a successful model in many circumstances, and it is still widely used nowadays. However, with the increasing studies of various ion channels, more types of P_o_–*V* relations were found that were hardly fitted by the Boltzmann equation. The reason is apparent that the Boltzmann model applies to a two-state process, containing only the open and closed states. But many ion-permeation processes can involve multiple open states or mix with inactivation states, which are not described by a two-state model.

In addition, the unitary single-channel conductance sometimes is not constant. An obvious case is the inward-rectifier potassium (Kir) channel that often shows two levels of single-channel conductance, for example, Yang and Jiang found that Kir4.1 channel in symmetrical 150 mM K^+^ solutions showed an inward conductance of 22 pS but a nearly zero outward conductance (5). Similar behaviors were also found in other channels, e.g., Moroni et al. found that the mechanosensitive Piezo1 channel in symmetrical Na^+^ solutions showed an inward conductance of 41.3 pS but an outward conductance of 27.1 pS (6). These behaviors are conveniently described by two levels of conductance that need be incorporated into a model.

To accurately represent the macroscopic ionic current, we present an improved model that incorporates the adapted single-channel conductance associated with the channel populations comprised of at least two states. With these modifications, the proposed model captures the essential features of several current-voltage relations and represents the current curves more accurately. We demonstrate the applicability of the proposed model by comparing model curves with the published experimental results.

## Methods

Intuitively, ionic current can be described by

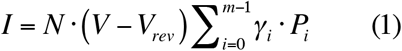

where *V* is the membrane potential, *V*_rev_ is the reversal potential, *N* is the number of channels examined. Suppose that the ionic current obtained over a range of voltage *V* involves *m* channel states ranged from 0 to *m*-1, and *P_i_* is the probability of the channel state *i, γ_i_* is the unitary conductance associated with the channel state *i*. Among these *m* states, we assume that there is a ground state with probability *P*_0_, e.g., a closed state that does not conduct ions so that *γ*_0_ = 0. If only two states are involved and *γ*_0_ = 0, the open probability P_o_ = *P*_1_, and the current I = *N*P_1_*γ*_1_(*V* – *V*_rev_), which reduces to the conventionally accepted current formula. Note that the proposed model does not necessarily require *γ*_0_ = 0, but in the ion-channel problem, there usually exists a nonconducting state represented by the ground state with *γ*_0_ = 0. Now according to the thermodynamic principles (7), at steady state:

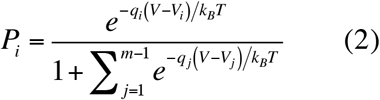

Where *q_i_* is the electric-charge change of the ion-protein complex in state *i* relative to the ground state, and *V_i_* is the half-activation potential associated with each state *i*. Note that this probability distribution also applies to the single channel behavior, because the ion permeation through a single channel can be decomposed into the steps of ion-binding then permeating through the channel. And according to the kinetic model (7), at steady state, the probability of each state can be described by Eq. 2.

Now considering the rectification behavior of ion channels, we model the macroscopic currents using separate values for the single-channel conductance related to the inward and outward currents, respectively:

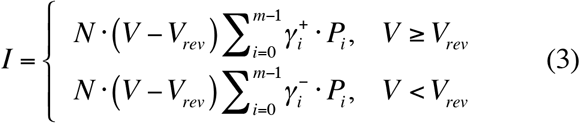

Here 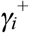 is one level of the *i*th-state single-channel conductance when *V* ≥ *V*_rev_, and 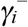 is the other level conductance of the same state when *V* < *V*_rev_. Note that 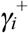 and 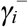 can differ or equal one another, when 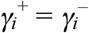, Eq. 3 reduces to Eq. 1.

In this paper, we examine the lower-rank models with *m* = 2, 3, and 4 only, because they are sufficient to describe the current-voltage curves of the selected cases. In other problems, the number of states *m* depends on the experimental data obtained from the single-channel recordings. In case of the two-state model, assuming that *γ*_0_ = 0 for the closed or the nonconducting ground-channel state,

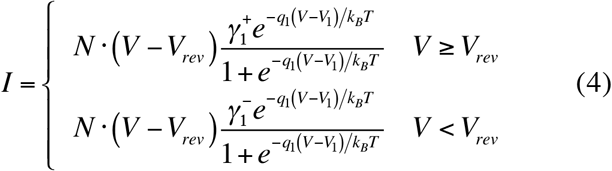

For the three-state model, still assuming that *γ*_0_ = 0 for the ground-channel state,

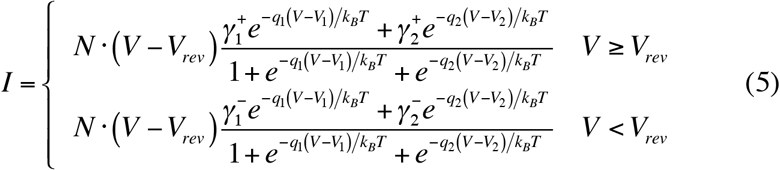

For the four-state model, when *γ*_0_ = 0,

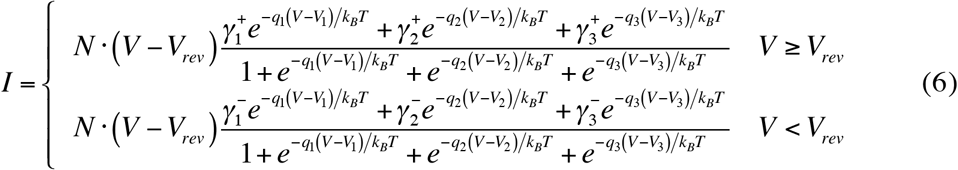

In Results and Discussion sections, the current-voltage relations were fitted assuming a zero ground-state conductance (*γ*_0_ = 0), and at least one open state with the probability *P*_1_. In the two-state model, P_o_ = *P*_1_, and the P_o_–*V* relation simply reduces to the Boltzmann equation. In the three-state and four-state models, 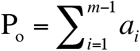 where *a_i_* = 1 for *γ_i_* ≠ 0, and *a_i_* = 0 for *γ_i_* = 0. In these cases, *P_i_* is modeled by Eq. 2 with *m* = 3 or 4.

The experimental data used in this paper were obtained directly from the published papers (as presented in their Results sections, with some data converted from the curves fitted by the Boltzmann equation in the original papers). Here, using the three-state or four-state models, the data were initially fitted by the non-linear least-squares method, then adjusted manually so as to match the single-channel experimental results. All model parameters used in this paper were presented in the supplemental Tables 1-3. Channel structures were rendered by Chimera (8).

## Results

In the patch-clamp experiments, single-channel recordings are often used to elucidate the ion-permeation mechanisms. In these experiments, P_o_ can be calculated by several methods, especially the variance-analysis method (9), and *γ* can be obtained from the relation of the single-channel current amplitude and the test voltage (represented by the *i–V* curve in this paper). Thus, some of the model parameters are directly obtained from the experiments, leaving fewer ambiguities when fitting the remaining parameters. This greatly improves the quality of modeling the ionic currents.

### Description of the two-state model

We do not elaborate on the two-state case when *γ*_1_^+^ = *γ*_1_^−^, since this is already widely used and commonly referred to as Boltzmann equation. Here we only show the altered I–*V* curves when *γ*_1_^+^ ≠ *γ*_1_^−^. Suppose that an ion channel opens at the depolarizing potentials and closes at the hyperpolarizing potentials such that *q*_1_ = −1 e and *V*_1_ = −20 mV. Simply setting *N* = 100 and *γ*_1_^+^ = *γ*_1_^−^ pS, the single-channel current amplitude *i–V* curve, the open probability P_o_–*V* curve, and the macroscopic current I–*V* curves all are obtained, and are shown in Fig. 1A. Now keeping *γ*_1_^+^ unchanged and varying *γ*_1_^−^ to 10 or 50 pS, the I–*V* curve shifts downward or upward in the inward current; whereas keeping *γ*_1_^−^ unchanged and varying *γ*_1_^+^ to 10 or 50 pS, the I–*V* curve bends down or up in the outward current (Fig. 1A). Thus, the unequal values of *γ*_1_^+^ and *γ*_1_^−^ can produce different effects on the I–*V* curves.

**Figure 1.**
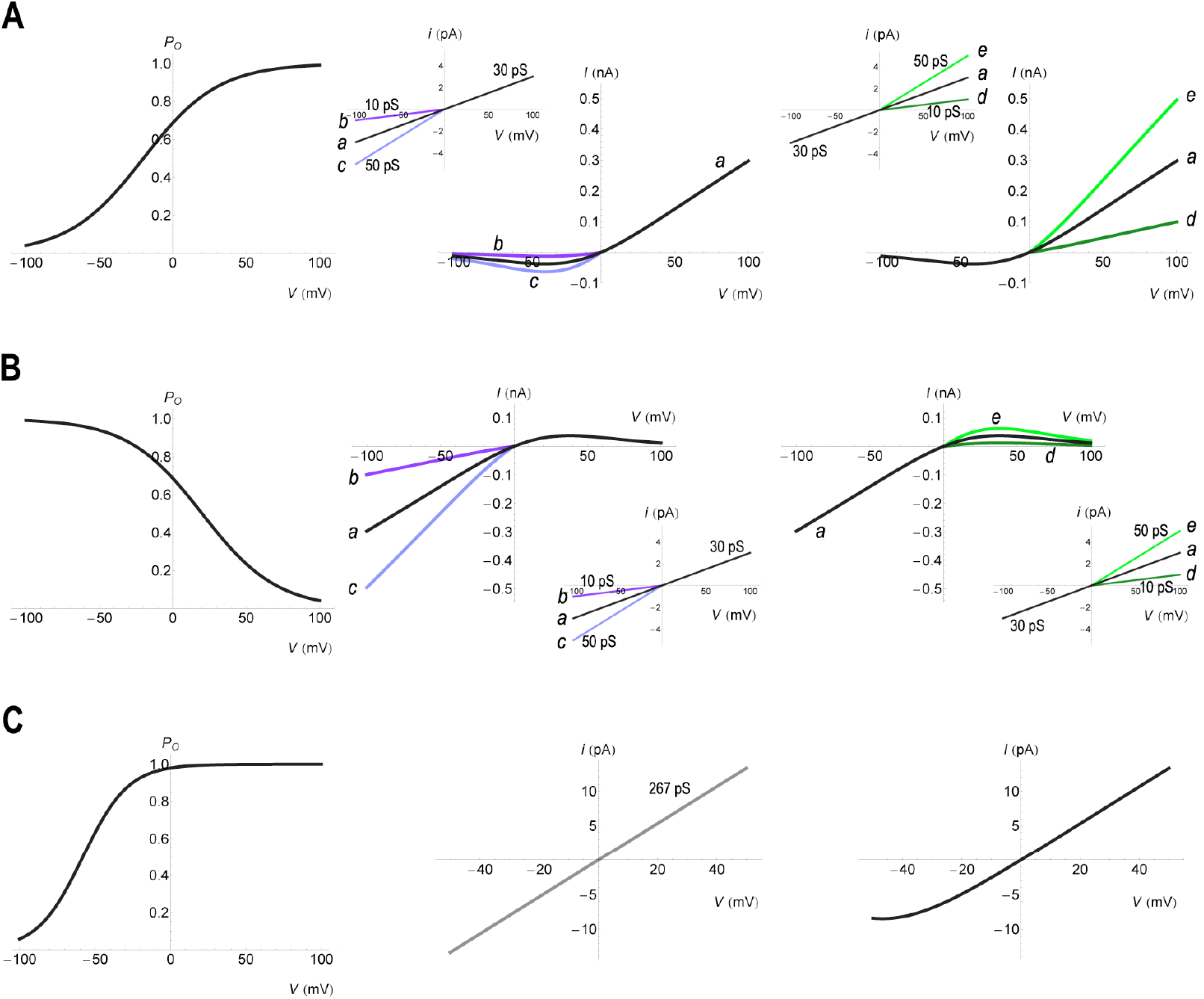
Two-state model curves. **A**, Depolarization-induced channel opening curves (*q*_1_ = −1 e, *V*_1_ = −20 mV). The open probability P_o_–*V* curves are same for all cases *a-e*. The single-channel current amplitude *i–V* curves differ in the values of *g*_1_^−^ (10 pS for case *b*, 50 pS for case *c*) or *g*_1_^+^ (10 pS for case *d*, 50 pS for case *e*), compared with case *a* (*g*_1_^+^ = *g*_1_^−^ = 30 pS). The changes of conductance were reflected in the macroscopic current I–*V* curves. **B**, Hyperpolarization-induced channel opening curves (*q*_1_ = 1 e, *V*_1_ = 20 mV). P_o_–*V* curves are same for cases *a-e*. *i*–*V* curves differ in values of *g*_1_^−^ (10 pS for case *b*, 50 pS for case *c*) or *g*_1_^+^ (10 pS for case *d*, 50 pS for case *e*), compared with case *a* (*g*_1_^+^ = *g*_1_^−^ = 30 pS). **C**, BK channel model curves (*q*_1_ = −1.77 e, *V*_1_ = −58 mV). Above −30 mV, BK channel almost fully opens that exhibits nearly an ohmic behavior. All model parameters are listed in Supplemental Table 1.

Similarly, the diagonally symmetric current curves are obtained (Fig. 1B) if the channel opens at hyperpolarizing potentials such that *q*_1_ = 1 e and *V*_1_ = 20 mV. Varying the model parameters as above, we see the corresponding changes in the macroscopic currents analogous to those shown in Fig. 1A.

There is a special case called the ohmic behavior, which is represented by a linear relation in I–*V* curve. This can occur when the channel fully opens over the test voltages. For example, Van Welie and Du Lac found that under 1 μM intracellular Ca^2+^ concentration and near physiological temperature (10), the large-conductance Ca^2+^-activated potassium (BK) channel showed a unitary single-channel conductance (*γ*_1_^+^ = *γ*_1_^−^ = 267 pS), and it opened at negative potentials with the half-activation potential *V*_1_ = −58 mV and a calculated *q*_1_ = −1.77 e (converted from the slope factor of 15 mV in Ref (10)). According to Eq. 4, this means that the BK channel almost fully opens above −30 mV that shows nearly an ohmic behavior in the I–*V* curve (current above −30 mV in Fig. 1C).

### Modeling I–*V* curve of a BK channel

BK channel (11-13) adheres to an allosteric gating scheme (13-18). It is activated by the intracellular Ca^2+^ ions that bind to the cytoplasmic gating ring composed of RCK1 and RCK2 (regulators of K^+^ conductance) subunits to regulate gating (19–24). Meanwhile it contains the voltage-sensing domain (VSD) that responds to the voltage changes (13, 25, 26). Structural studies show that the voltage sensor likely interacts with the RCK1 N-lobe via the S4-S5 linker (16), fulfilling the controls of allosteric gating. In addition, residues in the deep-pore region may also participate in gating because mutations of deep-pore residues can greatly influence the channel currents (27–30). Besides these, BK channel can coassemble with the auxiliary *β*1-4 subunits (31–33) or *γ* subunits (34, 35), which regulate or influence gating as well (36–42). The P_o_–*V* curve often (but not always) shows a two-state sigmoidal shape well characterized by the Boltzmann equation (13, 25, 43), where *V*_1_ shifts to the more hyperpolarized potentials with the elevated intracellular Ca^2+^ concentrations.

Among these results, Wu et al. reported a P_o_–*V* curve of a BK channel activated by 1 mM intracellular Ca^2+^ that showed a gradually reducing P_o_ towards the depolarizing potentials (44). The P_o_–*V* curve was reconstructed here by the three-state model with one open state that showed *q*_1_ = −1.44 e, *V*_1_ = −24 mV, *q*_2_ = −2.71 e, and *V*_2_ = 42 mV (Fig. 2A). Compared with the parameters (44) of fitting the left part of the curve using the Boltzmann equation: *q*_1_ = −1.44 e, *V*_1_ = −24.2 mV, the three-state model shows an extra state with *γ*_2_ = 0 developing quickly at positive voltages. Note that the experimental P_o_–*V* curve was obtained by dividing the averaged ramp current by the single channel current curve. Thus, such P_o_–*V* curve can also be described by the three-state model containing two open states with one of the open states having a smaller conductance that develops towards the depolarizing potentials. However, we exclude this choice because Wu et al. reported a unitary conductance of this BK channel under the symmetrical K^+^ solution, which was 235 pS (44). Therefore, it’s reasonable that this BK channel opens from a single state with the unitary conductance *γ*_1_^+^ = *γ*_1_^−^ = 235 pS.

**Figure 2.**
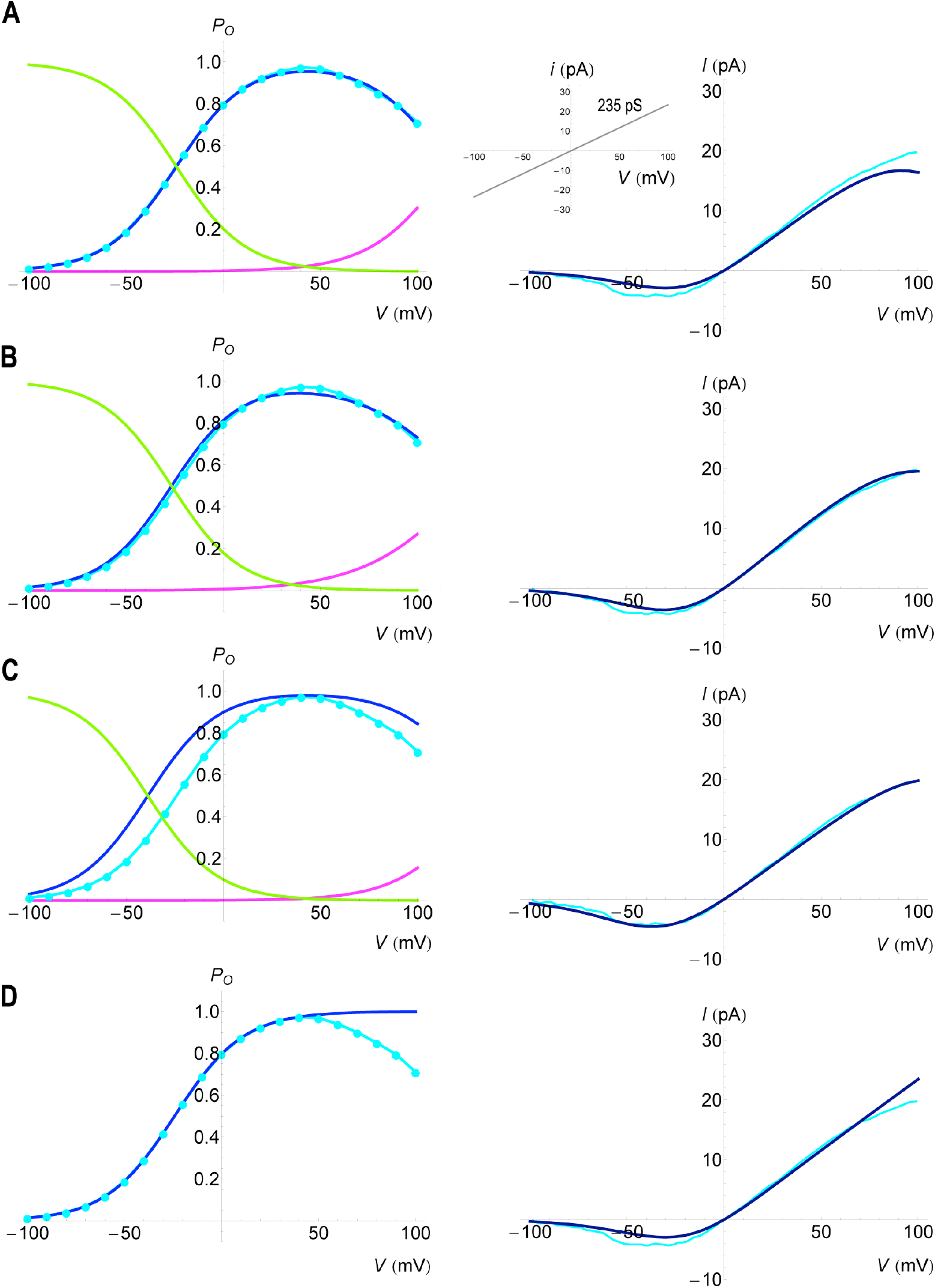
Model curves for a BK channel in symmetrical 115 mM KCl solutions. The experimental P_o_–*V* and I–*V* curves are colored in cyan. Model I–*V* curves are colored in dark blue. In P_o_–*V* curves, all three-state models contain one open state (state 1 in blue, *g*_1_^+^ = *g*_1_^−^ = 235 pS), one closed state (state 0 in green, *g*_0_^+^ = *g*_0_^−^ = 0 pS), and one nonconducting state (state 2 in magenta, *g*_2_^+^ = *g*_2_^−^ = 0 pS). For comparison, parameters for the three-state models are: **A**, *q*_1_ = −1.44 e, *V*_1_ = −24 mV, *q*_2_ = −2.71 e, *V*_2_ = 42 mV, and *N* = 1. **B**, *q*_1_ = −1.44 e, *V*_1_ = −26.5 mV, *q*_2_ = −2.41 e, *V*_2_ = 35 mV, and *N* = 1.14. **C**, *q*_1_ = −1.44 e, *V*_1_ = −39 mV, *q*_2_ = −2.71 e, *V*_2_ = 42 mV, and *N* = 1. Parameters for the two-state model are: **D**, *q*_1_ = −1.44 e, *V*_1_ = −24.2 mV, and *N* = 1. The left part of the experimental P_o_–*V* curve follows the Boltzmann equation of parameters used in **D**. All three-state and two-state models use the same *i*–*V* curve. And all model parameters are listed in Supplemental Table 2.

With these model parameters, we obtain the I–*V* curve that resembles but deviates a little from the experimental result (Fig. 2A). Apparently, the model curve underestimates the experimental I–*V* curve. They match better if *N* = 1.14 with slightly modified *V*_1_, *q*_2_, and *V*_2_ values (Fig. 2B). But the experimental I–*V* curve was the average of 20 measurements in a single patch that likely contained only one channel, whereas the P_o_–*V* curve was calculated from the averaged ramp currents of 3~5 patches (44). Therefore, *N* = 1 is preferred. With *N* = 1, the I–*V* curve can match the experimental result if the P_o_–*V* curve is slightly modified (Fig. 2C).

Interestingly, this modified P_o_–*V* curve was obtained simply by shifting *V*_1_ (from −24 mV in Fig. 2A to −39 mV in Fig. 2C). This means that the single channel opens at a slightly more negative potential (around −39 mV) when measuring the I–*V* curve. This is reasonable because a single channel often opens stochastically that the P_o_–*V* and I–*V* curves measured in different patches or at different time periods may show hysteresis. Hence these model curves show the impact of one parameter (shifted *V*_1_) to the variation of the macroscopic current.

In fact, several well-known allosteric gating models (14, 15, 45, 46), and some block models representing the inactivation imposed by the auxiliary *β*2 or *β*3 subunits (39, 47, 48) were already proposed for BK channels. Hence more than three channel states are likely involved in several circumstances, especially when BK channel is regulated by the auxiliary subunits (39, 47, 48) or blocked by inhibitors (49–51). But here, for this specific problem, the BK channel exhibits a constant single-channel conductance over the test voltages, that likely involves only three channel states, or at most three major states over the test voltages examined. The contributions from the minor states (if they exist) will be small compared with those of the major states. Thus the three-state model having one open state with conductance *γ*_1_^+^ = *γ*_1_^−^ = 235 pS can describe this ion-permeation process. Although simplified, it captures the essential channel features necessary to describe the mechanism, and it shows better representation of the ionic current than that obtained by the Boltzmann equation (Fig. 2D).

### Modeling I–*V* curve of a MthK channel

The MthK K^+^ channel is a voltage-dependent Ca^2+^-activated K^+^ channel, resembling the BK channel (52–54). Although the voltage sensors are missing, the gating ring at the intracellular side is preserved that senses Ca^2+^ concentration to regulate gating just like the BK channel (52, 55-57). In addition, the selectivity filter of MthK channel may also play important roles in regulating ion permeation (58-61). Besides these functional domains, the N-terminal region can affect channel inactivation as well (62). Recently, Fan et al. reported the MthK channel’s cryo-EM structure where the N-terminus plugged into the channel cavity, blocking the ion-permeation pathway via the ball-and-chain mechanism (54). Several well-designed models were already developed for MthK channel, describing the macroscopic currents blocked by inhibitors, which clearly showed that the channel functions relied on more than two states (63, 64).

In one experiment, Thomson and Rothberg (65) showed the P_o_–*V* relation (cyan curve in Fig. 3A) of K^+^ ions permeating through the MthK channel in 200 mM symmetrical K^+^ solutions. The MthK channels were activated by 20 mM intracellular Ca^2+^, then blocked by Ca^2+^ ions at high depolarizing voltages (53, 58). At hyperpolarizing voltages, P_o_ decreased due to the flickering ion-permeation events (65). Apparently, this is not a two-state P_o_–*V* curve.

**Figure 3.**
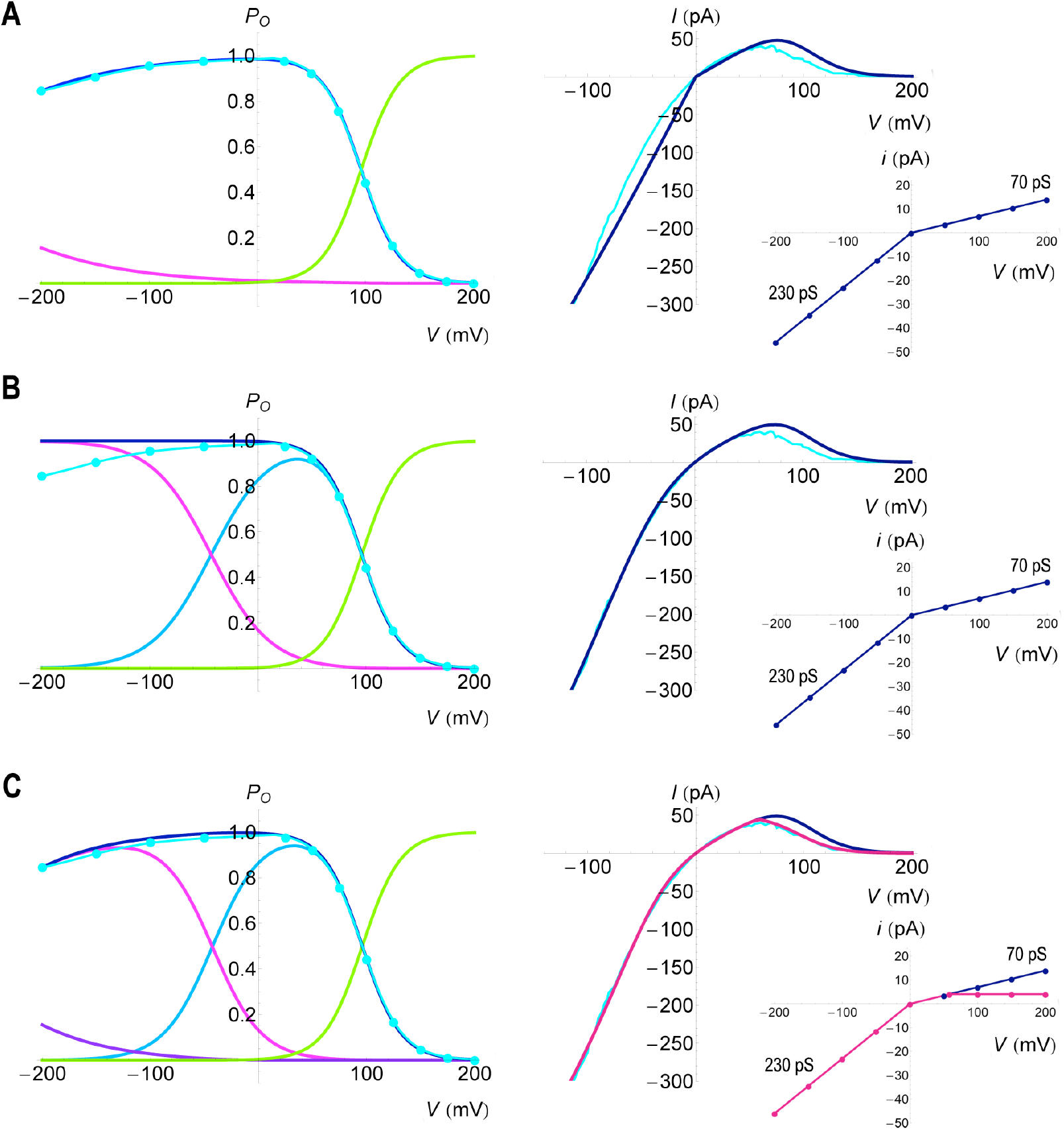
Model curves for MthK channels in symmetrical 200 mM KCl solutions. The experimental P_o_–*V* and I–*V* curves are colored in cyan. Model I–*V* curves are in dark blue and pink. **A**, Three-state model with one open state (state 1 in blue, *g*_1_^+^ = 70 pS and *g*_1_^−^ = 230 pS) and two nonconducting states (state 0 in green with *g*_0_^+^ = *g*_0_^−^ = 0 pS, and state 2 in magenta with *g*_2_^+^ = *g*_2_^−^ = 0 pS). **B**, Three-state model with two open states (state 1 in blue with *g*_1_^+^ = *g*_1_^−^ = 70 pS, and state 2 in magenta with *g*_2_^+^ = *g*_2_^−^ = 230 pS) and one nonconducting state (state 0 in green with *g*_0_^+^ = *g*_0_^−^ = 0 pS). **C**, Four-state model with two open states (state 1 in blue with *g*_1_^+^ = *g*_1_^−^ = 70 pS, and state 2 in magenta with *g*_2_^+^ = *g*_2_^−^ = 230 pS) and two nonconducting states (state 0 in green with *g*_0_^+^ = *g*_0_^−^ = 0 pS, and state 3 in purple with *g*_3_^+^ = *g*_3_^−^ = 0 pS). In **C**, the outward current is modified by an adaptive *i*–*V* curve (pink), so that the model fits the I–*V* curve better (pink). See text for details of the modification. The right part of the experimental P_o_–*V* curve follows the Boltzmann equation of parameters *q*_1_ = 1.4 e and *V*_1_ = 96 mV. All model P_o_–*V* curves are rendered in dark-blue color. And all model parameters are listed in Supplemental Table 3.

First, we model this process using the three-state model. The single-channel recording experiments clearly showed that there were at least two levels of the single-channel conductance associated with the inward and outward currents, respectively (65). Zadek and Nimigean had reported in a previous study that the MthK channel activated by 5 mM Ca^2+^ showed an inward conductance of 242 pS, and an outward conductance of 27 pS at 150 mV (equivalent to 81 pS at 50 mV if assuming an unchanged current above 50 mV in accordance with their result plotted in Fig. 1C) (53). Here, using one open state with two levels of conductance (*γ*_1_^+^ = 70 pS and *γ*_1_^−^ = 230 pS based on Fig. 1A of Ref. (65)) and two nonconducting states (one develops at the positive potentials with *γ*_0_^+^ = *γ*_0_^-^ = 0 pS and another develops at the negative potentials with *γ*_2_^+^ = *γ*_2_^−^ = 0 pS), the P_o_–*V* curve matches the experimental curve, but the I–*V* curve does not (Fig. 3A).

Then we fit the curve using two open states (with *γ*_1_^+^ = *γ*_1_^−^ = 70 pS and *γ*_2_^+^ = *γ*_2_^−^ = 230 pS) and one nonconducting state (*γ*_0_^+^ = *γ*_0_^−^ = 0 pS). This time the I–*V* curve matches better, but the P_o_–*V* curve simply shows a sigmoidal behavior that fully opens at hyperpolarizing potentials without any reductions (Fig. 3B). Clearly, it requires another inactivation state at negative potentials.

So we fit the curve using the four-state model with two open states (with *γ*_1_^+^ = *γ*_1_^−^ = 70 pS and *γ*_2_^+^ = *γ*_2_^−^ = 230 pS) and two nonconducting states (*γ*_0_^+^ = *γ*_0_^−^ = 0 pS and *γ*_3_^+^ = *γ*_3_^−^ = 0 pS). This time, both P_o_–*V* and I–*V* curves resemble the experimental results, but the I–*V* curve deviates slightly at positive potentials (dark-blue curve in Fig. 3C), possibly due to the saturation current through single channels mixed with the ion-block effects as described by Cox et al. while studying a BK channel current (66). Here we model this outward current of MthK channel by setting *γ*_1_^+^ = 70 pS for *V* < 58 mV, and *γ*_1_^+^ = 70 · (*V*_u_ – *V*_rev_)/(*V* – *V*_rev_) pS for 58 mV ≤ *V* < 200 mV, where *V*_u_ = 58 mV. This modification keeps the current amplitude constant above 58 mV, resembling the experimental results where the current amplitude changed little at 50 mV, 100 mV, 150 mV, and 200 mV (65). With this modification, both P_o_–*V* (dark blue) and I–*V* (pink) curves match the experimental results (cyan curves in Fig. 3C). And the calculated *N* = 12, which is also similar to the experimental result (around 10 single channels) (65).

Thus, this four-state model shows that K^+^ ions permeating through the MthK channel can involve two open states, one dominates at the positive potentials with a small conductance of 70 pS and one develops at negative potentials with a large conductance of 230 pS. An inactivation state develops quickly at positive potentials, whereas another inactivation state develops slowly at negative potentials that possibly relates to the flickering gating events (65). Examining the model curves, the overall P_o_–*V* curve is clearly decomposed into contributions of two open states having different levels of conductance (Fig. 3C).

## Discussion

The proposed model was developed based on the patch-clamp data. Thus, the number of sub-channel states *m* (not chosen randomly) must match the single-channel recording data and it depends on the single-channel current curve and the unitary conductance obtained from the experiments. Hence *m*, *V*_rev_, *γ*_i_^+^, and *γ*_i_^−^ values all are obtained from the experiments, and the only model parameters left to find are *q_i_* and *V_i_* of each sub-channel state, and the number of channels *N*. Therefore, this model can be used to predict the number of channels in the patch, to quantify I–*V* relations so as to compare the different channel functions (such as comparing *q_i_* and *V_i_* among different channels, or estimating the populations of the sub-channel states at particular voltages using Eq. 2), and most importantly, the model helps uncover the hidden mechanisms common to several ion channels that are particularly useful to study their structure-function relations. Here is an example.

MthK channel resembles BK channel in many aspects (52–54) but differs in the macroscopic currents. Both channels comprise S5 and S6 transmembrane helices containing the pore region, and a large intracellular gating ring sensing the Ca^2+^ concentration which, upon elevation, shifts the activation potential of both channels to the hyperpolarizing voltages (13, 25, 58). However, BK channel usually displays an outward-rectifying current whereas MthK channel shows an inward-rectifying current. Apparently, these are due to their markedly different P_o_–*V* and *i*–*V* curves, which are often used to describe their ion-permeation mechanisms. Does this mean that the two channels employ different mechanisms despite bearing the high structural resemblance?

Here, the model helps reveal the common mechanism that is not seen in two channels’ overall P_o_–*V* curves, which appear quite differently in the figure (cyan curves in Fig. 2C and Fig. 3C). Resemblance appears when the model curves are plotted together (Fig. 4A): the sub-states of two channels demonstrate comparable populations distributed over the voltage axis. These sub-state populations contain similar shapes, shifting along the voltage axis reflecting their different activation potentials that may be partly related to the different Ca^2+^ concentrations used in the two experiments. And it is convenient to compare these curves using the model parameters. For example, we can compare the parameters of the relative sub-state populations described by the green and blue curves of two channels (Fig. 4A): for BK channel, it is described by the population ratio of state 1 to state 0, i.e., *P*_1_/*P*_0_ = exp(−*q*_1_(*V* – *V*_1_)/*k_B_T*); analogously, for MthK channel, it is described by the population ratio of state 1 to state 2, i.e.,

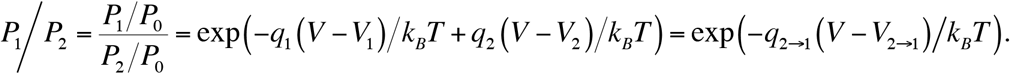

*q*_2→1_ and *V*_2→1_ are easily calculated when *q*_1_, *V*_1_, *q*_2_, and *V*_2_ are known. Thus *q*_1_ = −1.44 e and *V*_1_ = −39 mV for BK channel are compared with *q*_2→1_ = −1.13 e and *V*_2→1_ = −42.73 mV for MthK channel for these curves. Similarly, the relative populations described by the magenta and blue curves (Fig. 4A) can be compared by the parameters *q*_2→1_ = 1.27 e and *V*_2→1_ = 133.84 mV (for BK channel) with *q*_1_ = 1.44 e and *V*_1_ = 96 mV (for MthK channel).

**Figure 4.**
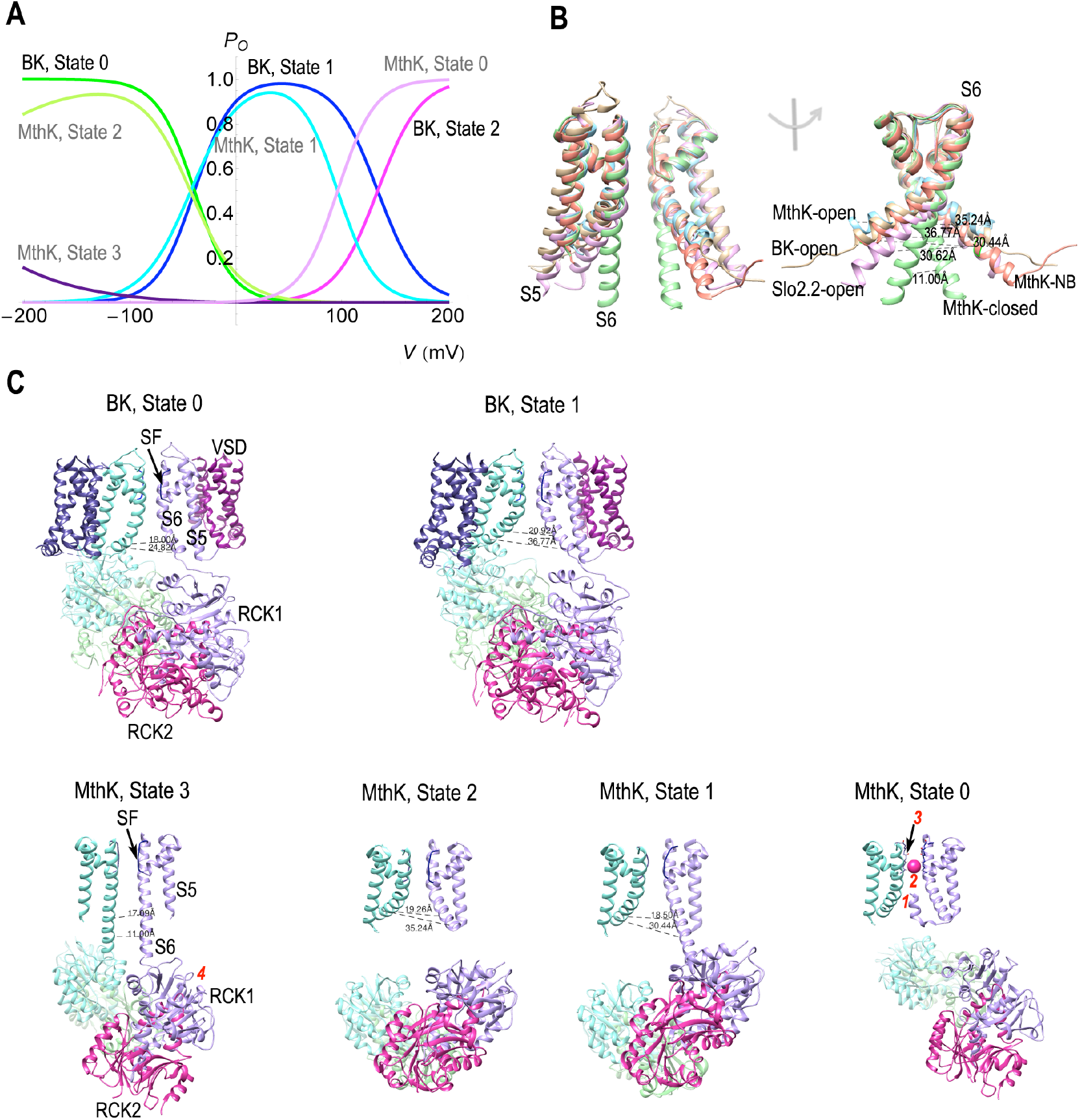
State-dependent structural resemblance of BK and MthK channels. **A**, *P_i_–V* curves of the BK (Fig. 2C) and MthK (Fig. 3C) channel substates are plotted together. **B**, Structural alignments of an open BK channel (yellow, PDB 5tj6), an open MthK channel (light blue, PDB 6ux7), an open Slo2.2 channel (purple, PDB 5u70), a closed MthK channel (green, PDB 6u5r), and an N-terminus blocked MthK channel (MthK-NB, orange, PDB 6u68, the N-terminus plugged into the cavity is not shown). For clarity, only two subunits containing the S5 and S6 transmembrane helices are shown on the left, and only S6 helices are shown on the right. These structures differ in the opening scales at the inner gate regulated by the swing of S6-helix tails, represented here by the different diameter values between the Cα carbons of residues (G316 for BK, I99 for MthK, and M333 for Slo2.2 channels) on the opposite subunits, which are 36.77 Å, 35.24 Å, 30.62 Å, 11 Å, and 30.44 Å for the channels mentioned above. **C**, Putatively corresponding structures for the sub-channel states described in **A**. BK and MthK channels both contain the S5-S6 transmembrance helices containing the selectivity filter (SF), and a large intracellular gating ring (RCK1 and RCK2, RCK2 is a separably free domain in MthK channel), but MthK channel lacks the voltagesensing domain (VSD). For clarity, only two subunits are shown, and the opening scale at the inner gate is described by two diameters of Cα carbons located at opposite subunits (P309 and G316 for BK channel, E92 and I99 for MthK channel). State 0 of BK channel (PDB 5tji) resembles state 3 of MthK channel (PDB 6u5r), but the inner gate of BK channel is relatively large, and it is unclear if this large gate persists under the hyperpolarizing potentials. The open structure of BK channel state 1 (PDB 5tj6) is comparable to the open structure of MthK channel state 2 (PDB 6ux7). The smallconductance MthK channel state 1 is putatively related to the small opening scale at the inner gate (although other possibilities may also apply here), analogous to the N-terminus blocked structure (PDB 6u68) but prior to the N-terminus plugged into the cavity, just like a preinactivated open state. The structure of BK channel state 2 (not shown) may resemble those of MthK channel state 0, which comprises at least four possible structures: 1, an N-terminus blocked structure (PDB 6u68); 2, a hypothesized ion-blocked structure; 3, a putatively inactivated filter structure (not shown here); 4, a closed MthK structure (resembling MthK channel state 3).

What can be inferred from these similar sub-state population curves? Realizing that structural changes can be state dependent, we speculate that the two channels may share a common mechanism involving similar sub-channel states which, upon stimuli (either voltage or ligand alterations), can undergo similar conformational switches at particular voltages. Currently, several well-described crystal or cryo-EM structures were published for Slo (Slo1, an alternative name for BK, and Slo2.2) (16, 67-69) and MthK channels (52, 54) that greatly improved our understanding of their functions. Here we try to relate these well-defined structures to each channel state so as to compare their structure-function relations in the same frame as described by Fig. 4A.

At first glance, the major open states of two channels (state 1) populated at positive voltages likely correlate with the Ca^2+^-bound open structures of Slo1 (69) and MthK (54) channels. But state 1 of BK channel has a large conductance of 235 pS, whereas that of MthK channel has a reduced conductance of 70 pS. Here we correlate state 1 of MthK channel to an N-terminus blocked structure (54), but prior to the N-terminus flipped into the channel cavity, just like a preinactivated state. This structure shows a reduced opening diameter at the inner gate compared with the fully opened MthK structure (Fig. 4B). Interestingly, an analogous structure (68) of a Na^+^-activated Slo2.2 channel also showed a small opening scale at the inner gate (Fig. 4B). And comparable to MthK channel, Slo2.2 channel also showed the inward-rectifying current in symmetrical K^+^ solutions (70) and a nonlinear *i*–*V* curve at positive voltages (71). The estimated conductance of Slo2.2 channel was around 100-180 pS, smaller than that estimated for Slo1 channel (100-270 pS) (72). Therefore, we speculate that the reduced conductance correlates with the small opening at the inner gate, just like the conclusion in a previous report by Naranjo et al. (73). Thus the Ca^2+^-activated fully open structure is more likely associated with state 2 of MthK channel that has a high conductance of 230 pS. Whereas state 1 of MthK channel can correlate to a preinactivated state with a reduced inner-gate diameter but prior to the N-terminus flipped into the channel cavity, analogous to the preinactivated open state described by Lingle et al. while studying the BK channel inactivated by the *β*3 subunit (39, 48). However, other possibilities such as a partially ion-blocked structure may also apply in this scenario (not shown in Fig. 4C).

State 0 of BK channel resembles state 2 of MthK channel in population (Fig. 4A), but one is nonconducting (*γ*_0_^−^ = 0 pS for BK channel) whereas another is highly conductive (*γ*_2_^−^ = 230 pS for MthK channel). The difference may be explained by the voltage-sensing domain that at negative potentials the voltage sensor of BK channel may switch to a conformation that tends to close the inner gate (possibly through interactions with the RCK1 N-lobes (16)) and renders the channel nonconductive, whereas the MthK channel can remain conductive due to the lack of the voltage sensors. Note that BK channel usually remains open under the high Ca^2+^ concentration, why does it switch to a closed structure in state 0? Structural studies find that even under the high agonist concentrations, a small portion of Slo2.2 or MthK channels can remain closed, e.g., 7% MthK channels having closed structures coexist with the open structures under 5 mM Ca^2+^ concentration (54), and 17% Na^+^-activated Slo2.2 channels remain in closed structures under 300 mM Na^+^ concentration (68). Therefore, state 0 of BK channel possibly bears a closed structure with a particular voltage-sensor conformation locked by the hyperpolarizing potentials (a hypothesis is that negative potentials may help increase the population of the closed structures). Similarly, the closed inner-gate structure of MthK channel (54) is possibly related to the state 3 of MthK channel (Fig. 4C).

At depolarizing voltages, the inactivation states of both channels contain similar shapes in population (Fig. 4A), which may result from the N-terminus block (54) or ion block (74) or some hypothesized nonconducting state of the selectivity filter (58-61). In addition, we cannot exclude the possibility of the closed channel structures for these inactivation states, because in some cases of the ion-blocking Slo2.2 channel, different ions may compete for the same allosteric sites that influence channel gating as well (70). Furthermore, in other circumstances, BK channel inactivation can be regulated by the coassembled auxiliary subunits, e.g., the *β*2 and *β*3 subunits can promote BK channel inactivation via the N-terminus of *β* subunits (36–38), similar to a ball-and-chain inactivation process. Thus there are several possibilities linked to the inactivation state of two channels developed at depolarizing potentials. Whether these possible (some are hypothesized) conformations can work alone or in a concerted manner or through sequential steps to induce inactivation requires further examinations.

The above comparison relates the published BK and MthK channel structures to each sub-channel state, based on the assumption that the two channels operate via the same mechanism. This comparison predicts that BK channel may switch to an inactivation state bearing a structure just like that of the MthK channel in state 0, although the inactivated BK channel structure has not been reported yet. In addition, this comparison predicts that the voltage-sensing domain may play important roles in shaping two channels’ ionic currents, that with it BK channel becomes nonconductive at negative potentials (state 0) showing only the outward-rectifying current, but without it MthK channel is highly conductive at the same voltage range (state 2) showing the inward-rectifying current.

It is well known that BK and MthK channel gating depends on both ligands and voltage (13-18). This allosteric gating in BK channel is regulated by the interplay between voltage sensors and RCK1 N-lobes (16), and hence ligands and voltage can produce similar effects in channel activation. Here, we have only examined the results of the high ligand-concentration activated currents, and we speculate that similar curves exist under the low-ligand concentrations, but those curves may shift along the voltage-axis reflecting their different activation potentials, and some sub-channel states may not appear over a limited voltage range examined. In addition, the BK channel regulated by auxiliary subunits or under other circumstances may also involve multiple open states, which may be incorporated into the model using the proper single-channel recording data.

Nevertheless, the main conclusion as outlined by Fig. 4A may still retain that BK and MthK channels can operate on the same working cycle and employ similar conformational switches. On this cycle, BK channel operates at positive voltages showing the outward-rectifying current, whereas MthK channel operates at negative voltages showing the inward-rectifying current owning to the persistent large-conductance open state 2 over the negative potentials.

## Supporting material

Supporting material includes three tables.

## Acknowledgments

This work was supported by Natural Science Foundation of China (Grant No. 21003023).

