## Supplemental Tables 1-3 for "Modeling Macroscopic Currents of Ion Channels"

**Supplemental Table 1.** *Two-state model parameters for curves plotted in Fig. 1.*

| Curve No. | $q_1$ (e) | $V_1$ (mV) | $g_1^+$ (pS) | $g_1^-$ (pS) | $V_{rev}$ (mV) | $N$ | $T$ (ºC) |
|---|---|---|---|---|---|---|---|
| 1A.*a* | -1 | -20 | 30 | 30 | 0 | 100 | 22 |
| 1A.*b* | -1 | -20 | 30 | 10 | 0 | 100 | 22 |
| 1A.*c* | -1 | -20 | 30 | 50 | 0 | 100 | 22 |
| 1A.*d* | -1 | -20 | 10 | 30 | 0 | 100 | 22 |
| 1A.*e* | -1 | -20 | 50 | 30 | 0 | 100 | 22 |
| 1B.*a* | 1 | 20 | 30 | 30 | 0 | 100 | 22 |
| 1B.*b* | 1 | 20 | 30 | 10 | 0 | 100 | 22 |
| 1B.*c* | 1 | 20 | 30 | 50 | 0 | 100 | 22 |
| 1B.*d* | 1 | 20 | 10 | 30 | 0 | 100 | 22 |
| 1B.*e* | 1 | 20 | 50 | 30 | 0 | 100 | 22 |
| 1C | -1.77 | -58 | 267 | 267 | 0 | 1 | 34 |

**Supplemental Table 2.** *Model parameters for curves plotted in Fig. 2.*

| Curve No. | $q_1$ (e) | $V_1$ (mV) | $q_2$ (e) | $V_2$ (mV) | $g_1^+ = g_1^-$ (pS) | $g_2^+ = g_2^-$ (pS) | $V_{rev}$ (mV) | $N$ | $T$ (ºC) |
|---|---|---|---|---|---|---|---|---|---|
| 2A[a] | -1.44 | -24 | -2.71 | 42 | 235 | 0 | 0 | 1 | 22 |
| 2B[b] | -1.44 | -26.5 | -2.41 | 35 | 235 | 0 | 0 | 1.14 | 22 |
| 2C[c] | -1.44 | -39 | -2.71 | 42 | 235 | 0 | 0 | 1 | 22 |
| 2D[d] | -1.44 | -24.2 | / | / | 235 | / | 0 | 1 | 22 |

[a,b,c]Three-state model.
[d]Two-state model.

**Supplemental Table 3.** *Model parameters for curves plotted in Fig. 3.*

| Curve No. | $q_1$ (e) | $V_1$ (mV) | $q_2$ (e) | $V_2$ (mV) | $q_3$ (e) | $V_3$ (mV) | $g_1^+$ (pS) | $g_1^-$ (pS) | $g_2$ (pS)[d] | $g_3$ (pS)[e] | $V_{rev}$ (mV) | $N$ | $T$ (ºC) |
|---|---|---|---|---|---|---|---|---|---|---|---|---|---|
| 3A[a] | 1.44 | 96 | 1.79 | 14 | / | / | 70 | 230 | 0 | / | 0 | 12 | 23 |
| 3B[b] | 1.44 | 96 | 2.38 | 41 | / | / | 70 | 70 | 230 | / | 0 | 12 | 23 |
| 3C[c] | 1.44 | 96 | 2.57 | 35 | 3.03 | -15 | 70 | 70 | 230 | 0 | 0 | 12 | 23 |

[a,b]Three-state model.
[c]Four-state model.
[d]$g_2 = g_2^+ = g_2^-$.
[e]$g_3 = g_3^+ = g_3^-$.
